# Vispro improves imaging analysis for Visium spatial transcriptomics

**DOI:** 10.1101/2024.10.07.617088

**Authors:** Huifang Ma, Yilong Qu, Anru R. Zhang, Zhicheng Ji

## Abstract

Spatial transcriptomics (ST) enables the comprehensive analysis of gene expression while preserving the spatial context of tissues. The histological images accompanying ST data provide spatially cohesive information that is often challenging to capture through gene expression alone. However, analyzing such images is challenging due to the presence of fiducial markers and background regions, which can obscure important features and complicate downstream analysis. To address these challenges, we developed Vispro, an end-to-end, fully automated image processing tool tailored for 10x Visium data. Vispro integrates four key modules of fiducial marker detection, fiducial marker removal and image restoration, tissue region detection, and segmentation of disconnected tissue areas. We demonstrated that Vispro enhances the quality of ST images and improves the performance of downstream analyses, including tissue segmentation, cell segmentation, image registration, and histology-based gene imputation.

## Introductions

Spatial transcriptomics (ST) is a transformative technology that simultaneously measures gene expression profiles, spatial locations, and imaging information from the same set of cells. Unlike single-cell sequencing, which cannot capture spatial or imaging data, ST provides a deeper understanding of how different cell types are spatially organized within a tissue and enables the study of cell morphology.

The imaging information accompanying ST has become indispensable in many analytical methods designed for ST data^1–11^, with two primary approaches currently being explored. One approach^1–9^ leverages image data to predict gene expression or facilitate gene analysis, aiming to extract molecular insights from histological images. A notable example is TESLA^9^, which estimates tissue contours and generates superpixel regions based on histological similarity, enabling high-resolution gene imputation.The other approach^10,11^ integrates both gene expression data and imaging information to develop more comprehensive models, using the spatial correlation between these two modalities to improve the accuracy and depth of biological interpretation. For example, SpaGCN^10^ integrates histological images with gene expression data and spatial cell locations to infer spatial domains and identify spatially variable genes. Similarly, stLearn^11^ combines imaging, gene expression, and spatial distances between cells to map spatiotemporal trajectories and cell–cell interactions. Additionally, imaging data has demonstrated significant potential in enhancing the accuracy of aligning multiple tissue sections, as highlighted by several studies^12^. This alignment facilitates integrative analysis of ST data across different samples, providing deeper insights into spatial gene expression patterns and tissue architecture.

Despite the promise of image-based approaches in ST, multiple challenges arise when analyzing images from ST datasets. First, images from 10x Visium, one of the most widely used ST platforms, include artificial reference points known as fiducial markers (Figure 1a,b), which are intended to outline tissue regions. However, these markers often create challenges during image processing. They can overlap with tissue regions due to operational errors, obscuring important structures and textures. Their blob-like shape frequently leads to misclassification as tissue structures, and their consistent square arrangement can confuse models attempting to learn spatial tissue patterns, leading to misinterpretation of global tissue architecture. Second, most ST images contain both tissue regions, where cellular structures are visible, and blank regions, referred to as background (Figure 1a,c). The background often contains residual stains and random noise, blurring tissue boundaries and interfering with gene expression predictions when entire images are used. Third, a single ST image may capture multiple tissue samples, particularly when tissue microarrays (TMAs) are used, containing tissues from different spatial locations or individuals (Figure 1a,d). Without proper segmentation of these distinct regions, downstream analysis becomes significantly more complicated.

**Figure 1.**
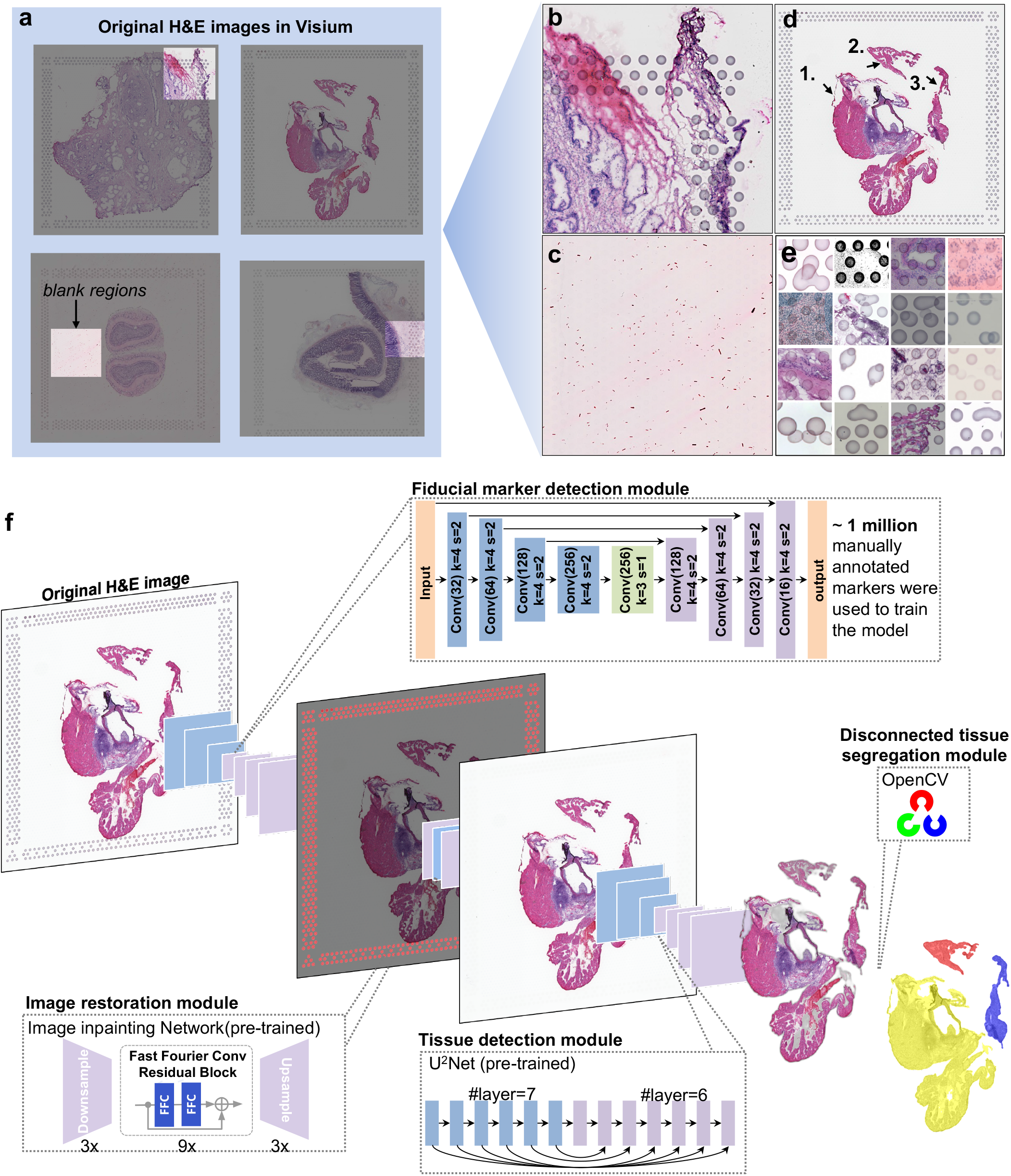
Vispro pipeline to improve the quality of Visium ST images. **a**, Original H&E-stained tissue images captured by the Visium platform. Zoomed-in views of the highlighted regions of interest are provided in **b-e. b**, An example showing fiducial markers overlaying tissue regions. **c**, An example showing background noise in the blank regions. **d**, An example of segregated tissue regions, with three subregions marked as 1, 2, and 3. **e**, Examples of different types of deformation in fiducial markers. **f**, The complete Vispro workflow, which consists of four modules: marker detection, image restoration, tissue detection, and disconnected tissue segregation.

These challenges have been largely overlooked and underexplored in the existing literature, particularly in relation to handling fiducial markers. The 10x Genomics Space Ranger^13^ and Loupe Browser^14^ can automatically or manually detect fiducial markers. However, they rely on a standardized fiducial marker template for alignment, assuming all markers are perfectly circular and uniformly arranged. This assumption fails to account for the deformations or misprints commonly present in real images (Figure 1a,e). More critically, these tools cannot automatically remove fiducial markers from images or recover distorted tissue regions where markers overlap with tissue. Additionally, existing methods for detecting and segmenting tissue regions are ineffective when applied directly to ST images without first removing fiducial markers, leading to erroneous results, as we will demonstrate. Although manual manipulation of images can address some of these issues, it is time-consuming and impractical for large-scale datasets. In summary, there is currently no method that can automatically and efficiently process ST images to fully remove technical artifacts and produce images ready for downstream analysis.

To address these challenges, we developed Vispro, a fully automated image processing tool for ST images (Figure 1f). Vispro includes four sequential modules: detecting fiducial markers, removing these markers and restoring the image, detecting tissue regions, and segregating disconnected tissue areas. The tool outputs cleaned images with accurately defined tissue regions, ready for integration into other image analysis software. We demonstrate that Vispro provides higher-quality image data in ST, including more accurate fiducial marker localization and tissue boundary detection. Additionally, we show that images processed by Vispro significantly improve the performance of downstream tasks such as cell segmentation, image registration, and gene imputation. In summary, Vispro is a valuable tool for optimizing the use of image data in ST.

## Results

### Vispro overview

Vispro is built upon a series of deep-learning models to perform image analysis tasks across its four modules (Figure 1f). The fiducial marker detection module employs a deep neural network based on U-Net^15^, with customized layers and loss functions designed to address the scale disparity between tiny fiducial markers and the large whole image. By emphasizing key shallow and deep layers and incorporating focal loss to address data imbalance, Vispro achieves improved accuracy in detecting small targets, outperforming the baseline U-Net. To train the model, we manually annotated approximately 100,000 fiducial markers from 167 ST images. The network produces a binary segmentation mask for fiducial markers. In the image restoration module, both the segmentation mask and the original image are fed into an image inpainting neural network, which restores the content in the marker area while preserving the surrounding image. The tissue detection module utilizes a U^2^Net neural network to identify the salient object and separate the tissue from the background. Finally, the disconnected tissue segregation module uses a brute-force search for pixel connectivity, implemented via the OpenCV library, to identify and segregate disjoint tissue regions. The complete pipeline operates in real time and in an end-to-end manner, with its efficiency detailed in Supplementary Table S1.

### Vispro accurately identifies and removes fiducial markers

Fiducial marker removal is not a simple task, as fiducial markers exhibit various levels of overlap with tissue regions in most ST datasets (Figure 2a). While a simple method such as image cropping can remove the fiducial markers, it also destroys parts of the tissue regions that may contain valuable information. A deep neural network, like the one employed by Vispro, is needed to effectively remove fiducial markers while preserving the integrity of the tissue regions.

**Figure 2.**
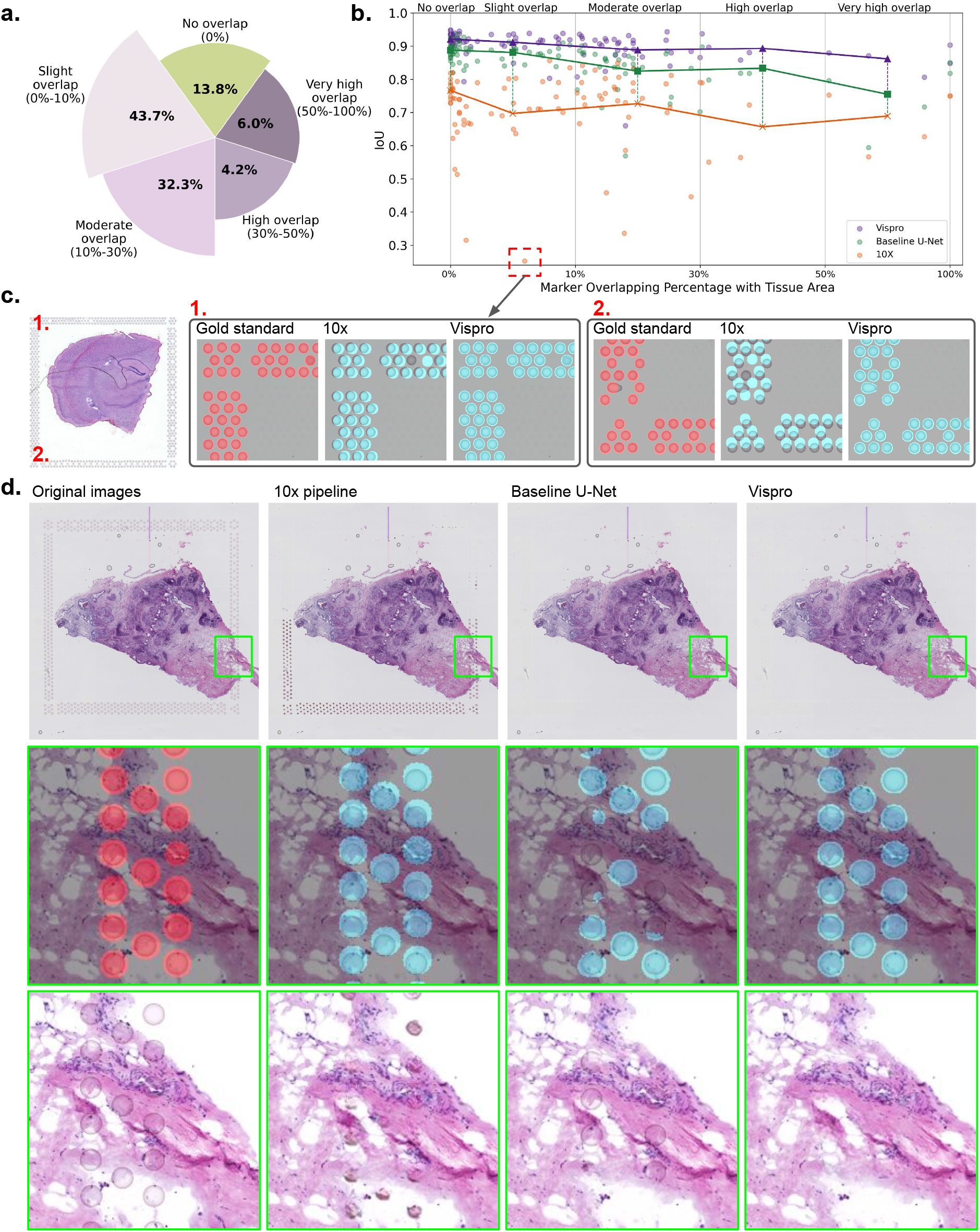
Fiducial marker removal and image restoration results. **a**, Proportion of images (number shown within each pie) with different overlap proportions (indicated outside each pie) between fiducial markers and tissue regions. The overlap proportion is calculated as the proportion of all fiducial markers that overlap with tissue regions and is indicated within parentheses in the text outside each pie. **b**, IoU of fiducial marker detection accuracy (y-axis) plotted against different overlap proportions (x-axis). Each point represents an image. **c**, An example Visium ST image showing zoomed-in views of the gold standard marker regions (red circles), fiducial markers detected by the 10x pipeline (cyan circles), and fiducial markers detected by Vispro (cyan circles), displayed from left to right in two distinct corners. **d**, A visual comparison of image restoration results. The three rows display, from top to bottom: the original images, gold standard fiducial marker locations (red circles) or fiducial markers identified by different methods (cyan circles) in a zoomed-in region, and the original image (first column) or the corresponding restored images (second to fourth columns).

We compared the performance of the 10x pipeline, the baseline U-Net architecture, and the fiducial marker detection module in Vispro for detecting fiducial markers in a cross-validation study (Methods). Figure 2b shows a quantitative comparison using the Intersection over Union (IoU) metric to evaluate the performance of all three methods against the manually annotated gold standard. Vispro achieves an IoU improvement of over 10% compared to the 10x Genomics pipeline and approximately 5% compared to the baseline U-Net across all levels of marker overlap, demonstrating its superior accuracy in identifying fiducial markers compared to existing methods. Notably, fiducial markers identified by the 10x pipeline exhibit significant misalignment with their actual locations in some cases. Figure 2c provides an example with an IoU below 0.3. In this example, the Visium slice shows global deformations, causing the fiducial marker arrangement to display shape distortions. Consequently, the 10x Genomics template struggles to align with the markers, resulting in missed fiducial markers (e.g., corner 1) and substantial displacement (e.g., corner 2). In contrast, the learning-based method employed by Vispro effectively leverages texture information to accurately localize the marker areas, with an IoU of 0.94.

Figure 2d illustrates the results of image restoration through inpainting the identified fiducial markers. The binary masks of fiducial markers obtained by the three methods were used as inputs to the same inpainting algorithm (Methods). Using the fiducial markers identified by Vispro, the inpainting process successfully removes all markers from both the background region and the tissue region in the images. In contrast, noticeable fiducial markers remain in the images when using the results from the 10x pipeline, while the baseline U-Net fails to recover some fiducial markers overlapping with the tissue region. This further demonstrates the superior accuracy of Vispro’s fiducial marker identification and underscores how the precision of fiducial marker detection significantly influences downstream image processing. Additional image restoration results are provided in Supplementary Figure S1.

### Vispro accurately identifies and segregates tissue regions

After inpainting the fiducial markers identified by Vispro, we evaluated Vispro’s performance in differentiating tissue regions from background regions (Figure 3) and in a subsequent module for segregating disconnected tissue regions (Figure 4). The computationally identified tissue regions and segregated tissue subregions were quantitatively compared to the gold standard of manually annotated tissue regions using IoU as the metric. For comparison, we also included results from the original images without the removal of fiducial markers.

**Figure 3.**
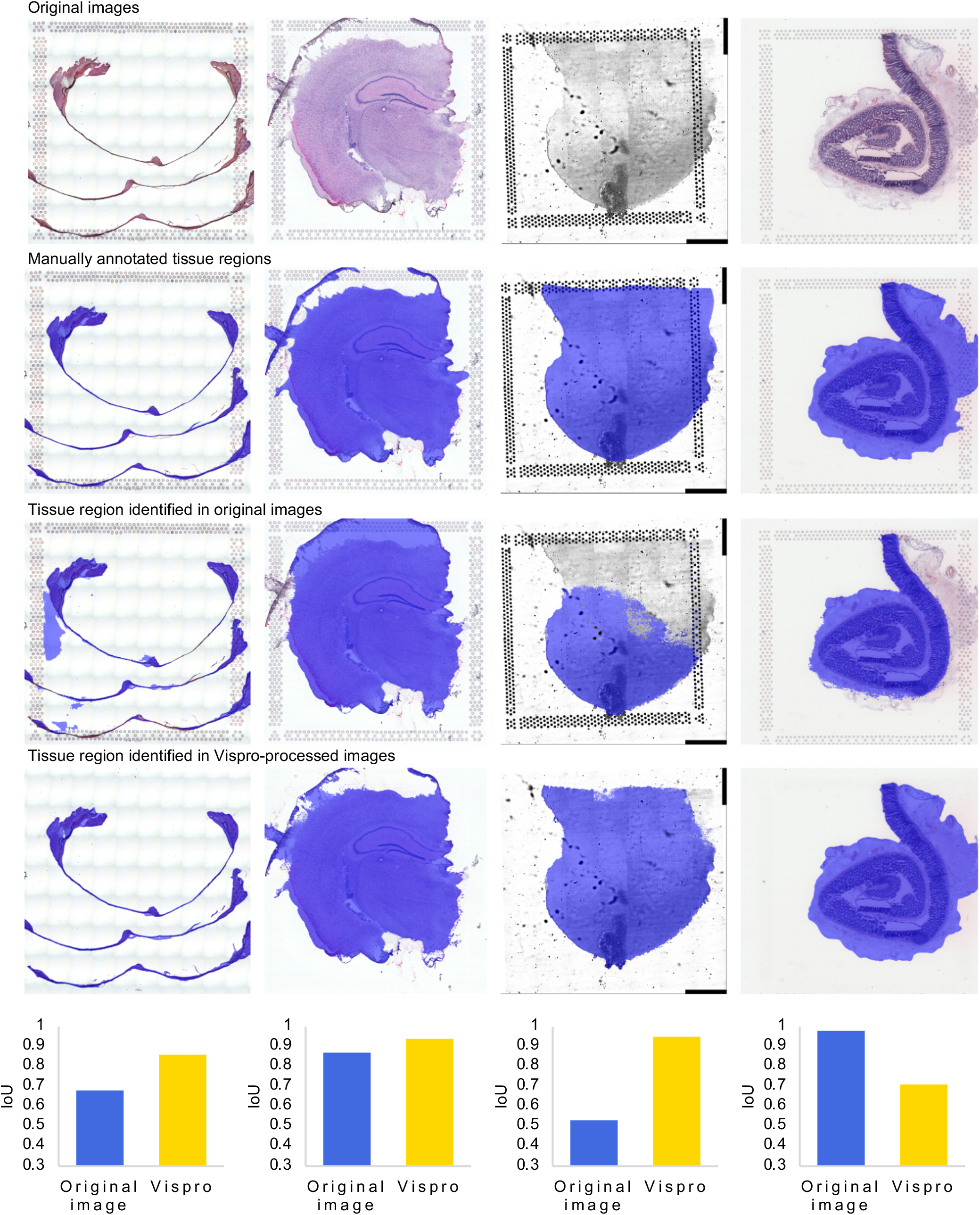
Tissue detection results. The five rows display, from top to bottom: the original images, the manually annotated tissue areas serving as the gold standard (blue regions), the detected tissue regions from the original image (blue regions), the detected tissue regions from the image after fiducial marker removal and image restoration by Vispro, and the IoU of tissue detection accuracy comparing the original image and the Vispro-processed image with the gold standard. Each column represents an image from a Visium sample.

**Figure 4.**
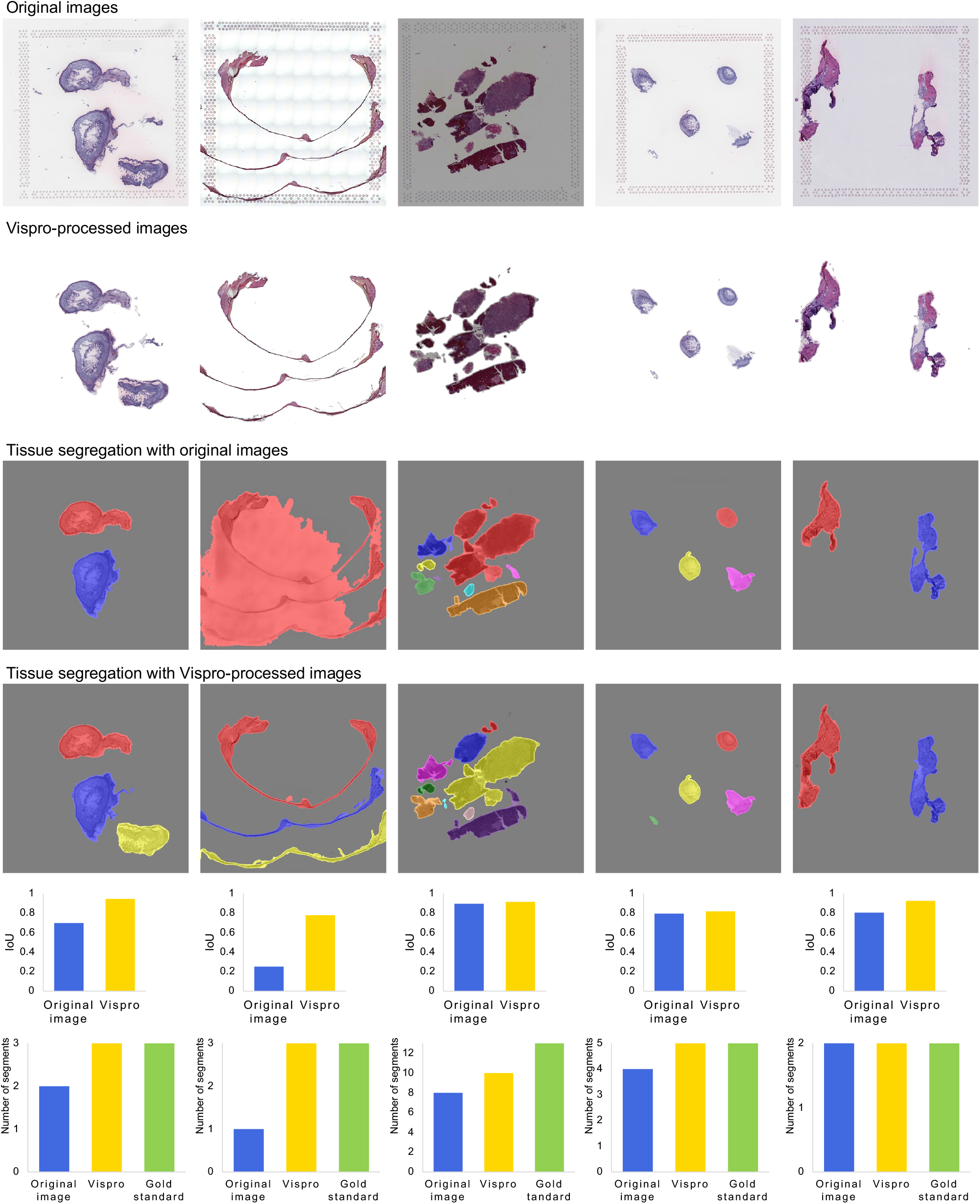
Disconnected tissue segregation results. The six rows display, from top to bottom: the original images, Vispro-processed images after tissue detection, tissue segregation results using the original image, tissue segregation results using the Vispro-processed image, IoU of tissue segregation accuracy comparing both images to the gold standard, and the number of segregated subregions comparing the original image, Vispro-processed image, and gold standard. Each column represents an image from a Visium sample.

Vispro accurately distinguishes tissue regions from background regions (Figure 3). The tissue regions identified by Vispro closely match the gold standard, with IoU metrics approaching 1. In contrast, directly identifying tissue regions without removing or inpainting the fiducial markers results in significantly worse outcomes, as evidenced by a notable drop in IoU. The presence of fiducial markers leads to enlarged or truncated tissue regions, particularly when the markers overlap with actual tissue areas.

After tissue region detection, Vispro effectively identifies disconnected tissue subregions (Figure 4). In most cases, it accurately detects the correct number of subregions. The IoU metric remains high, indicating that Vispro consistently predicts tissue areas with precision. In contrast, segmentation based solely on the original images shows a significant decline in performance, particularly by missing key tissue regions or incorrectly merging thin structures with the background, resulting in substantially lower IoU values.

### Vispro improves cell segmentation

Cell segmentation aims to identify and separate individual cells in an image^16,17^. Accurately identifying cell locations is crucial for understanding the spatial organization of different cell types within a tissue. Since most cell segmentation methods rely solely on imaging information, the quality of the images significantly impacts the performance of cell segmentation.

We applied StarDist^18^, a commonly used cell segmentation method, to three types of images: those processed by the full Vispro pipeline, images that only underwent the fiducial marker removal step of Vispro, and the original images (Figure 5). Since the goal of Vispro is to remove unwanted areas outside tissue regions, we only evaluated whether the cell segmentation method detected any cells in regions where no cells are expected, outside the tissue. These detections are considered false positives. The accuracy of cell segmentation within tissue regions is primarily dependent on the performance of the segmentation algorithm itself and is beyond the scope of this study.

**Figure 5.**
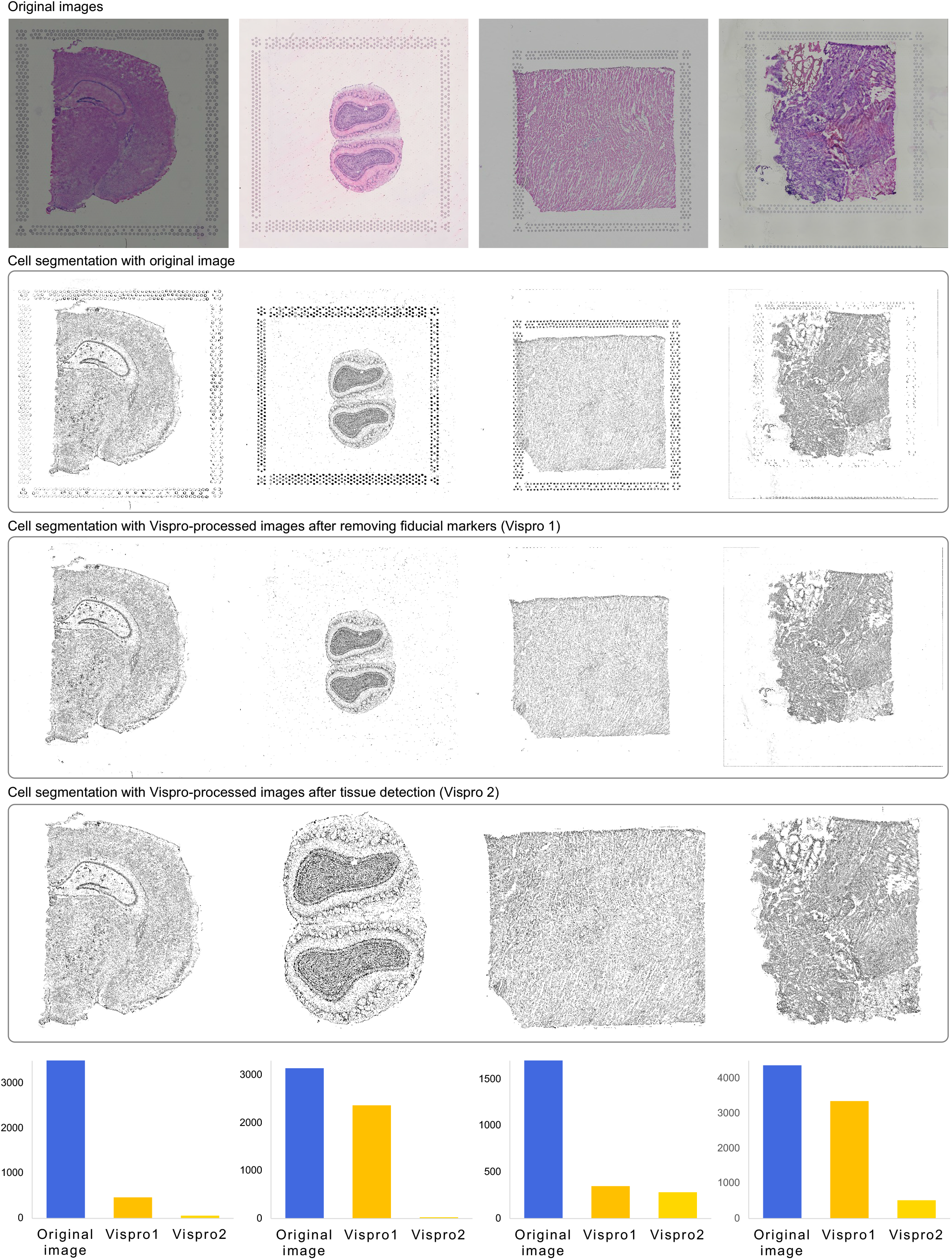
Cell segmentation results. The five rows display, from top to bottom: the original images, cell segmentation results using the original images, cell segmentation results with the Vispro-processed images after removing fiducial markers, cell segmentation results with the Vispro-processed images after tissue detection, and the number of segmented cells outside tissue regions. Each column represents an image from a Visium sample.

StarDist identifies thousands of false positive cells when applied to the original images or images that only underwent fiducial marker removal. This is primarily due to the segmentation method mistakenly identifying fiducial markers and background noise as cells, leading to numerous false positives. In contrast, the number of false positive cells is significantly reduced when using images processed with Vispro for segmentation, highlighting the importance of removing unwanted technical artifacts from the images. These findings suggest that Vispro enhances cell segmentation performance and leads to a more accurate characterization of the spatial locations of cells.

### Vispro improves image registration

Image registration involves transforming a source image into the same coordinate system as a target image so that the two images are directly comparable. Many studies generate multiple ST samples from similar tissue types^19^. Registering these images to a common coordinate system enables researchers to compare spatial and imaging features across individuals. Similar to cell segmentation, image registration relies solely on imaging data, and its performance is influenced by the quality of the images.

We tested the performance of bUnwarpJ^20^, a state-of-the-art image registration method, using images processed by the full Vispro pipeline and the original images (Figure 6). We considered two image registration settings. In the first setting (Figure 6a), both the source and target images were generated by the Visium workflow and included fiducial markers. In the second setting (Figure 6b), the source image was generated by the Visium workflow and included fiducial markers, while the target image was generated by the standard H&E histology workflow and did not include fiducial markers. This second setting is particularly useful for Visium CytAssist, an instrument that facilitates the transfer of transcriptomic probes from standard glass slides to Visium slides. The performance of image registration was quantitatively evaluated using two numerical measures—structural similarity index (SSIM) and mutual information (MI) (Methods).

**Figure 6.**
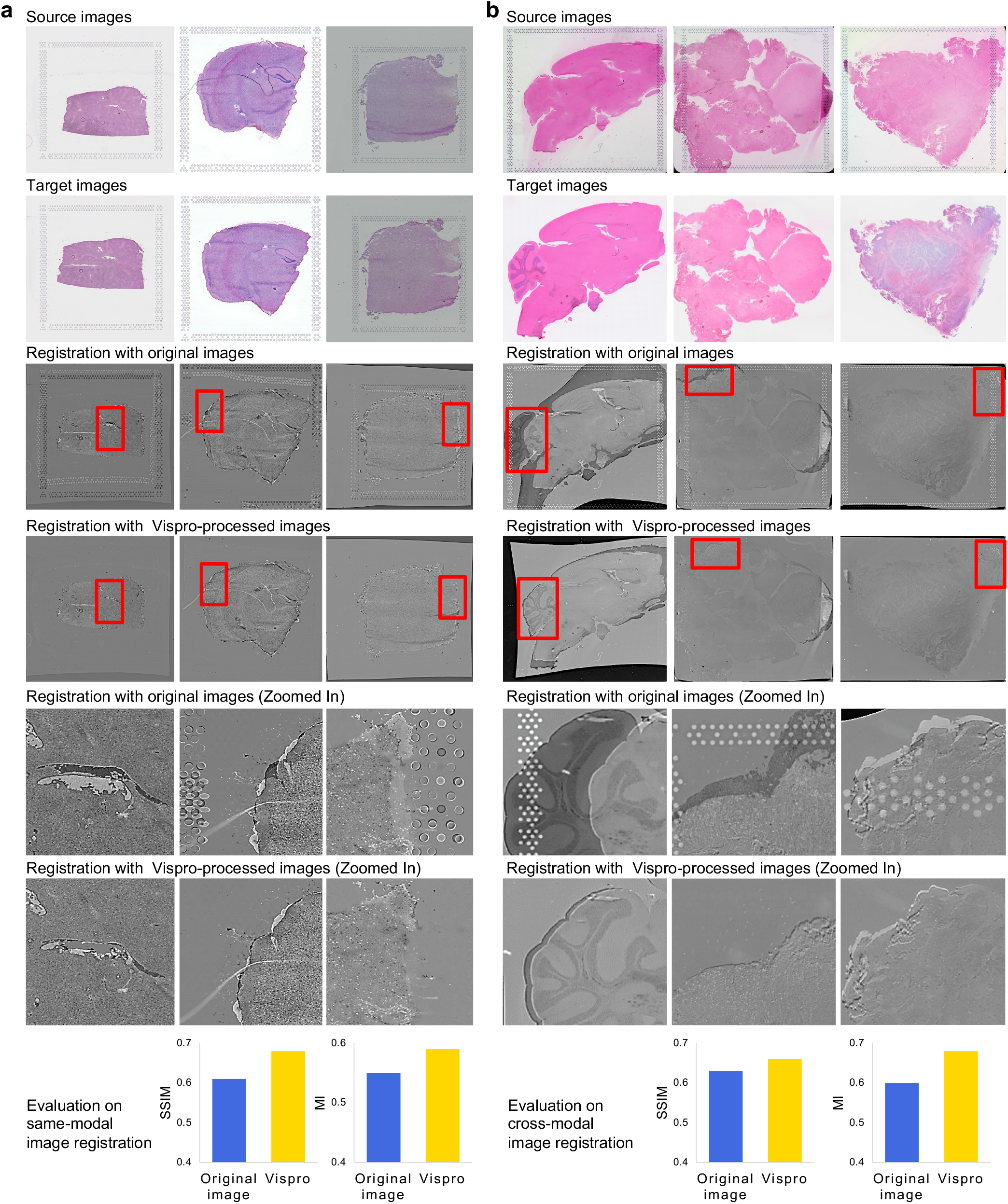
Image registration results. In **a**, both the source and target images were generated by Visium. In **b**, the source image was generated by Visium, while the target image was generated by the standard H&E workflow. The seven rows display, from top to bottom: the source images, the target images, image registration results using the original images, image registration results using Vispro-processed images, zoomed-in views of the results in the third row, zoomed-in views of the results in the fourth row, and the SSIM and MI metrics evaluating registration performance. Image registration results display the overlay of the warped source image and the target image in grayscale. The source image is presented with inverted intensity values, while the target image is shown with normal intensity values. Differences between the two images are highlighted as regions of high contrast, appearing as intensely white or black areas. Each column represents an image from a Visium sample.

Figure 6 demonstrates the performance of image registration in the two settings. For each case, a small section of the registered image was zoomed in to highlight the differences in registration results. When fiducial markers are present, bUnwarpJ attempts to align both the fiducial markers and the tissue regions. Since the position and layout of fiducial markers are influenced by technical factors (e.g., tissue placement on the slide), this results in inferior registration of some tissue regions, as seen in the misaligned tissue structures within the zoomed-in areas. In contrast, the performance of image registration improves substantially with images processed by Vispro. By removing fiducial markers, the registration process focuses solely on the tissue regions, resulting in better alignment of the highlighted tissue structures. Along with improved SSIM and MI values, these findings suggest that Vispro enhances the performance of image registration.

### Vispro improves histology-based gene imputation

Finally, we demonstrate how Vispro enhances the results of TESLA^9^, a method that imputes gene expression using histology image information. As examples, we examined the spatial expression patterns of senescence-associated genes, including CDKN1A, CDKN2A, and HMGB1^21,22^. While the original Visium data provides only discrete, spot-level spatial expression patterns for these genes, TESLA generates a continuous spatial gene expression map across the entire tissue.

We compared the results of TESLA with images processed by different pipelines as input (Figure 7). TESLA provides three options for tissue detection algorithms prior to performing gene imputation in tissue regions. However, none of these algorithms produce reliable results, causing TESLA to impute gene expression over large areas that do not correspond to actual tissue regions. This can lead to misleading interpretations of spatial gene expression patterns, particularly in regions not containing real tissue. In contrast, when using images processed by Vispro, TESLA’s imputation results closely align with the actual tissue regions, enabling more accurate interpretations of spatial gene expression patterns.

**Figure 7.**
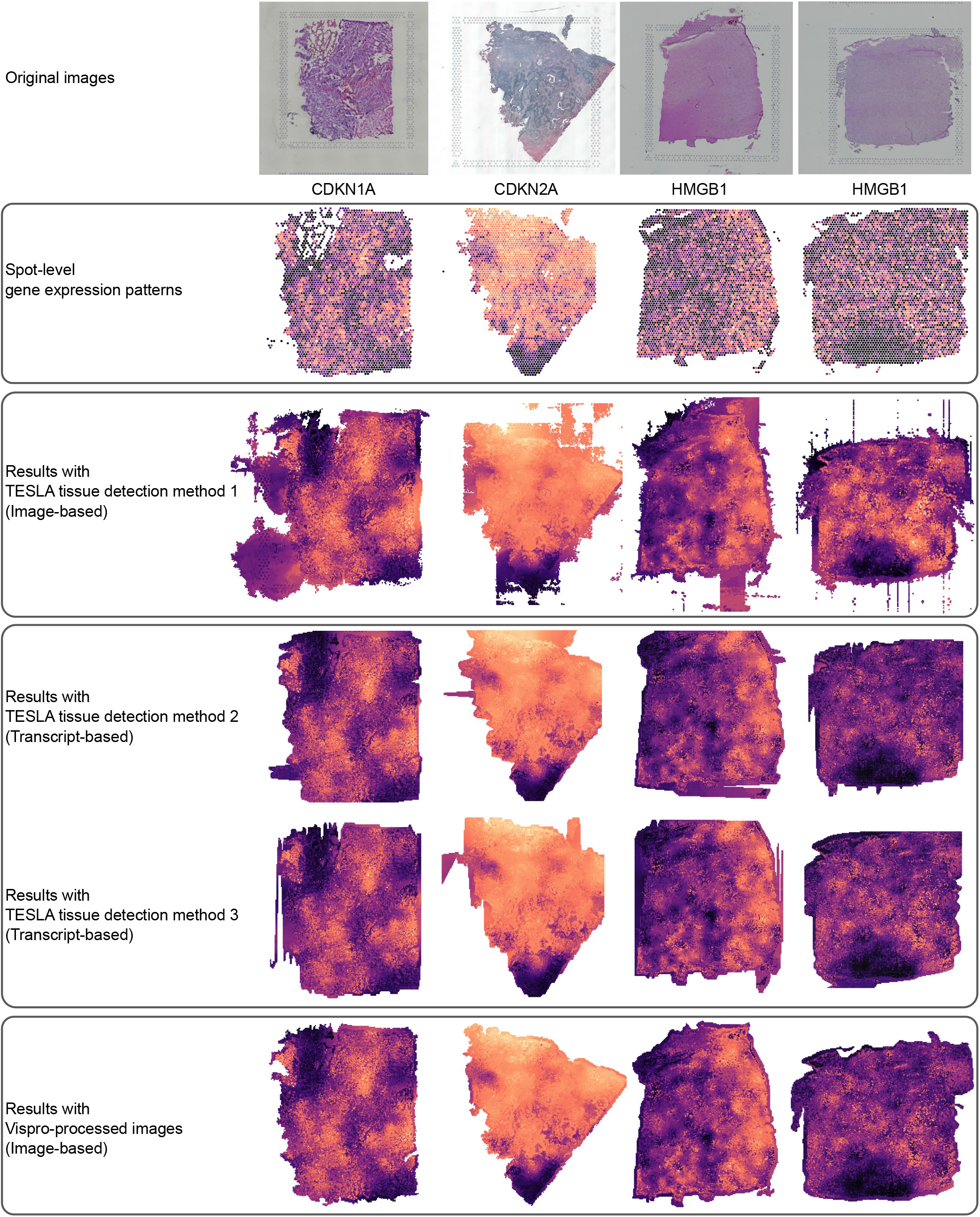
Image-based gene imputation results. The six rows display, from top to bottom: the original images, spot-level spatial gene expression patterns, TESLA results using TESLA’s first tissue detection method (canny contour detection), TESLA results using TESLA’s second tissue detection method (scanning transcript contour by spot x), TESLA results using TESLA’s third tissue detection method (scanning transcript contour by spot y), and TESLA results using Vispro-processed images. Each column represents an image from a Visium sample.

## Conclusion

We present Vispro, an automated image processing tool for ST images generated by the 10x Visium platform. Vispro provides four consecutive modules to clean ST images and extract images of tissue regions. It accurately identifies fiducial markers and tissue regions in real examples, outperforming the default processing pipeline provided by 10x. We have demonstrated that processing images with Vispro can substantially improve the performance of downstream imaging analysis tasks, including cell segmentation, image registration, and gene imputation.

## Methods

### Vispro module 1: Fiducial marker detection

The first step in Vispro involves detecting fiducial markers generated during the ST imaging process. Due to significant deformations in the shapes and positions of the markers, a deep learning approach was employed to capture their unique textures. We designed a neural network architecture based on the U-Net framework^15^ to address the specific characteristics of markers across entire tissue slices. Significant effort was dedicated to assembling a robust training dataset to support model development. The original H&E image is extremely large, making it impractical for network training. Therefore, the model was trained on high-resolution sub-images to localize the marker areas, which were subsequently projected back to the original image scale.

#### Training data collection

We collected 167 datasets generated by the 10x Visium platform from 48 studies, which were downloaded from STOmics DB^23^. The full list of studies is provided in Supplementary Table S2. To generate training annotations for fiducial marker areas, we applied the Hough Transform circle detection algorithm^24^, which uses pixel intensity to identify circular shapes in images. However, the algorithm produced numerous false positives in tissue regions and missed certain markers due to deformations or overlap with tissue areas. To address these limitations, we manually removed false positives and annotated markers missed by the Hough Transform using the Labelme data annotation tool^25^, resulting in a comprehensively annotated dataset containing 92,685 fiducial markers.

#### Neural network architecture

The neural network employed by Vispro is based on the U-Net architecture^15^, a widely recognized neural network design for medical image segmentation. U-Net features a symmetric encoder-decoder structure optimized for efficient feature representation and reconstruction. The encoder progressively extracts hierarchical features through downsampling, while the decoder restores spatial resolution via upsampling, enabling the integration of fine-grained details with contextual information. This fully convolutional architecture ensures seamless feature extraction and reconstruction, making it well-suited for segmentation tasks.

The implementation of Vispro incorporates four UNetDown blocks, each designed to progressively reduce spatial resolution by half while increasing the number of feature channels from the initial 3 (RGB) to 32, 64, 128, and 256, respectively. Each block uses a 4 × 4 convolution with a stride of 2, followed by instance normalization and ReLU activation^26^, forming the core structure of the UNetDown block.

To address the specific challenges of fiducial marker segmentation, characterized by small-scale structures that are globally distributed but constrained to a square shape, we introduced targeted modifications to the network design. These adjustments prioritize retaining fine-grained details in the initial encoding layers while enhancing global shape awareness in the deepest encoding layers. The network structure is detailed below:

The input of the network is a 3-channel image, i.e., *I* ∈ ℝ^3*×W×H*^.

For the first encoding block, additional convolutional layers are added while avoiding dropout to preserve detailed feature extraction.

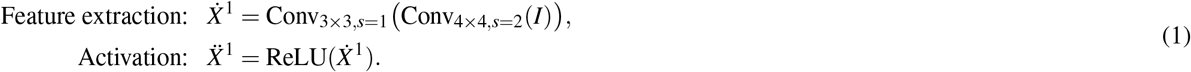

For the second and third encoding blocks (*i* = 2, 3), a foundation module is utilized.

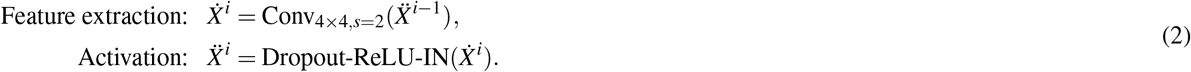

For the fourth encoding block, additional convolutional layers are added to enhance feature extraction.

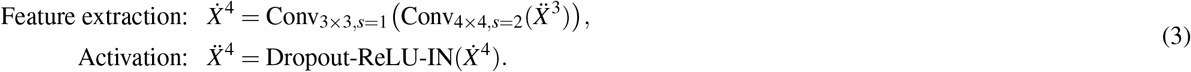

At the end of the encoding operations, additional bottleneck layers are incorporated at the deepest level of the network to further refine the receptive fields of the encoded features, as detailed below:

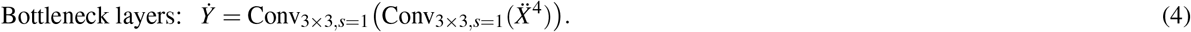

To reconstruct spatial context for fine-grained segmentation, the decoder progressively upsamples the bottleneck features through a series of four UNetUp blocks, mirroring the hierarchical structure of the encoder. At each scale, a 4 × 4 deconvolution operation is applied to double the spatial resolution of the learned features while sequentially reducing the feature channels from 256 to 128, 64, 32, and 16. These operations are followed by activation functions and skip connections, which seamlessly integrate high-resolution features from the encoder. This approach ensures the preservation of spatial details and enriches the reconstruction process with complementary contextual information.

Formally, the operations at the first decoding block are defined as follows, with additional convolutional layers applied to both the decoding features and the skip features to enhance global attention:

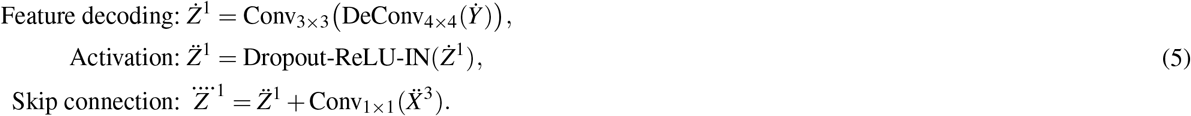

For the second decoding block:

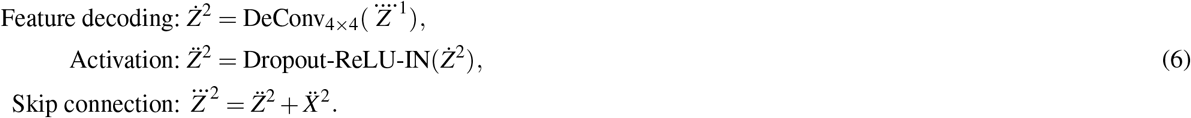

For the third decoding block:

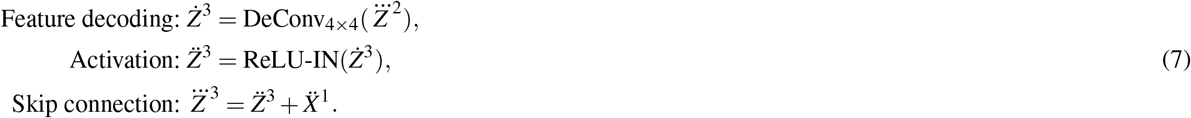

For the fourth decoding block:

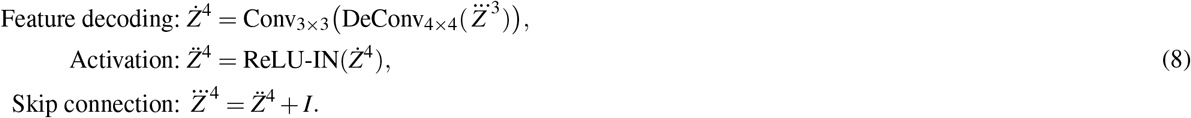

Each decoder layer processes the learned feature maps *Ż*^*i*^ while incorporating feature maps 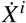 from the corresponding encoder layers via skip connections. These skip connections retain spatial details that are otherwise lost during downsampling, enabling the network to effectively merge high-resolution spatial features with deeper contextual information. This synergy significantly enhances the network’s capacity to reconstruct fine-grained features with precision, a critical advantage for segmentation tasks. The Vispro architecture further ensures that the network adeptly captures both intricate details and global context, making it particularly well-suited for segmenting small fiducial markers while maintaining their global shape constraints.

#### Neural network output

The output of the decoder is a 19-channel feature map at full resolution, comprising 16 learned feature channels and 3 channels from the original input image, i.e., 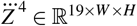. This feature map is passed through a learnable final layer, which generates the marker segmentation mask 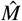:

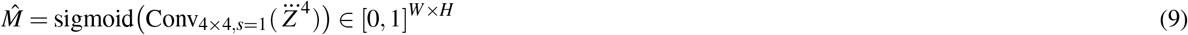

#### Loss function

We employed a combined loss function, consisting of Dice Loss^27^ and Focal Loss^28^, to evaluate the predicted marker mask, 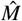, against the gold standard mask, *M*, obtained through manual annotation. The gold standard mask, *M*, is a binary mask where 1 represents the fiducial marker region and 0 represents the background region. In contrast, 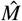 contains values in the range (0, 1), where values closer to 1 indicate a high likelihood of the marker region, and values closer 0 indicate a high likelihood of the background region.

The Dice Loss, widely used in segmentation tasks, evaluates the overlap between the target regions in the two masks. To calculate the Dice Loss, the predicted mask, 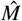, is first binarized by applying a threshold of 0.5, resulting in a binary prediction mask, 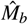. The Dice Loss is then computed as follows:

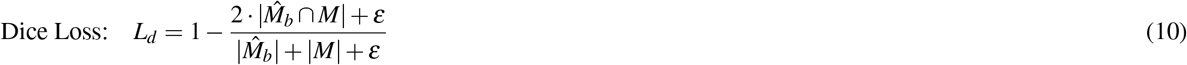

where *ε* is a smoothing factor set to *ε* = 1.0 to stabilize the loss value, especially for small or empty masks.

For the Focal Loss, it builds upon the pixel-wise Binary Cross-Entropy (BCE) Loss and introduces additional parameters to emphasize hard-to-classify markers while down-weighting easy-to-classify markers by adjusting the loss contribution of each example. Specifically, the BCE Loss is defined as:

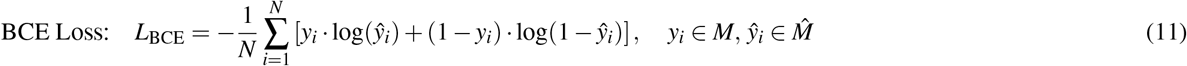

where *N* is the total number of pixels, *y*_*i*_ represents the ground truth label for pixel *i*, and *ŷ*_*i*_ is the corresponding predicted value from the neural network.

Focal Loss introduces weighting terms that dynamically adjust the loss contributions based on the predicted values of the network. This adjustment is achieved through the following formulation:

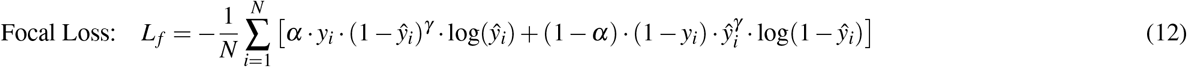

where *γ* controls the level of focus on hard examples, and *α* is a class-level weighting factor used to prioritize the target class. In Vispro, we set *α* = 0.95 and *γ* = 3.

By inversely scaling the weights based on the confidence levels of predictions, the loss function reduces the impact of well-classified samples while emphasizing harder and misclassified examples. This adaptive weighting mechanism ensures that the network prioritizes challenging samples during training, effectively balancing the contributions of both easy and hard examples to the overall loss.

The total loss used to train the network combines the Dice Loss and Focal Loss, defined as follows:

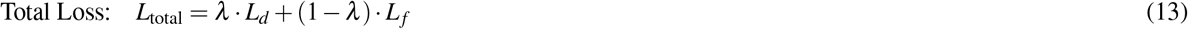

where *λ* is a weighting parameter that balances the contributions of the two loss components, set to *λ* = 0.9 in Vispro.

#### Training Procedure

Since markers located in the background region are more numerous and easier to classify, whereas markers overlapping with tissue regions are fewer and harder to distinguish, we implemented a sampling layer to increase the frequency of challenging samples during network training. Specifically, we computed a slice-wide marker overlap factor, defined as the ratio of in-tissue markers to the total number of markers. Based on this factor, we categorized images into three difficulty levels: hard examples (marker factor > 0.8), moderate examples (0.3 < marker factor ≤ 0.8), and easy examples (marker factor < 0.3). To ensure the model effectively learns from the most challenging cases, we increased the occurrence of hard samples threefold and moderate samples twofold in the training dataset.

During training, all images underwent carefully designed data augmentation strategies to enhance data variability and improve model robustness. These augmentations included random flipping (both left-right and up-down), scaling (0.8 to 1.0 times the original size), rotation (− 10^°^ to + 10^°^), brightness adjustment (0.5 to 1.2 times the original value), and Gaussian blurring (with a standard deviation randomly selected between 0 and 1). The model is trained using the Adam optimizer^29^ with a learning rate of 10^−4^. The optimizer’s parameters include *β*_1_ = 0.5 and *β*_2_ = 0.999, which control the exponential decay rates of the first- and second-order moment estimates of the gradients, respectively. These settings ensure stable convergence during training. The network was trained for 800 epochs.

### Vispro module 2: Image restoration

The second step in Vispro involves performing image restoration on the marker-covered regions of the images. For this task, we leverage LaMa, a deep learning model specifically designed for image inpainting^30^. Image inpainting is a computer vision technique that aims to fill in missing or corrupted regions of an image. The fundamental principle involves learning the surrounding textures and global image semantics to generate visually plausible content for the missing regions.

In this module, we incorporate the pre-trained model and parameters of LaMa^31^, which employs Fourier Convolutions to address the limitations of conventional inpainting methods that often struggle to generalize across varying resolutions. This approach ensures consistent performance on both low- and high-resolution images, enabling pre-training on large-scale natural image datasets while seamlessly adapting to ST images, which are typically high resolution.

The input to the model consists of the original image *I* and the predicted binary mask 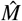 generated by Vispro Module 1. The output is an RGB image *Î*, where the fiducial marker regions are filled with plausible textures inferred from the surrounding image content.

### Vispro module 3: tissue detection

For the task of tissue detection, we employed the deep learning tool `backgroundremover`^32^, which is based on the U^2^-Net model^33^, a state-of-the-art salient object detection framework. The incorporation of Residual U-blocks (RSU) enhances the network’s capacity for efficient hierarchical feature learning, ensuring precise representation of fine-grained details alongside broader spatial patterns, making it particularly adept at robust tissue segmentation, especially in high-resolution images where traditional models frequently struggle to achieve a balance between detail preservation and global contextual accuracy.

By default, Vispro uses the `unet2` model and resizes images to 320 × 320 to optimize computational efficiency. For datasets with small tissue regions (less than 100 × 100 pixels), Vispro increases the image size to 1000 × 1000 and switches to the `u2netp` model. This adjustment enhances detection accuracy for finer structures and thinner tissue regions.

The input to the model is the restored RGB image *Î* produced by Vispro Module 2, and the outputs are a tissue mask 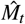 with values in the range [0, 1] and a blended RGBA image *Î*_*t*_ that visually highlights the detected tissue regions.

### Vispro module 4: disconnected tissue segregation

We identified disconnected components in the tissue probability mask 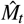 using the OpenCV library. Specifically, the probability mask was binarized using a tissue threshold value (default set to 0.8). The `label` function was then employed to count and label the disconnected regions in the binary mask. This function performs a brute-force search to determine pixel connectivity, iterating through each pixel in the mask and checking for 8-connectivity (including horizontal, vertical, and diagonal neighbors) to ascertain whether adjacent pixels belong to the same component. Once a connected group of pixels is identified, the algorithm assigns a unique integer label to the corresponding component. The output is a labeled mask in which each connected component is represented by a distinct integer value.

The input to this module is the tissue probability mask 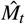 produced by Vispro Module 3, and the output is a refined segmentation mask 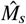, represented with integer values.

To improve the quality of the labeled mask, an additional step was introduced to remove small components, often artifacts caused by staining errors or computational inaccuracies near the tissue boundary. These small components were filtered out using a preset area threshold (default set to 20). Any component with an area below this threshold was excluded from the labeled mask. This refinement step ensures that the labeled mask more accurately represents tissue regions and minimizes noise.

### Competing methods

#### 10x standard processing pipeline

In the 10x pipeline, fiducial markers are processed using Space Ranger and Loupe Browser, which generate a layered image. This image includes a fiducial frame template with red circles overlaid on the original H&E image to highlight fiducial marker areas. These tools can automatically detect and highlight the fiducial regions, providing a visual reference for their locations on the tissue. However, they usually require complex model inputs, including gene count data, slide version information, and images. Additionally, in cases where fiducial markers are obstructed or tissue boundaries are unclear, manual alignment is often necessary.

To integrate 10x’s aligenment result into our workflow for evaluation, we first perform an RGB thresholding operation on the layered image to isolate the red color component representing the fiducial markers. This step extracts the red circles by identifying pixels that fall within a predefined range of RGB color intensities (R >180, G<30, and B<30). Once the red components are isolated, we apply a circle detection of Hough Circle Transform to detect the circular fiducial markers. This algorithm scans the image for circular shapes by identifying edge pixels and fitting circles to the detected contours. After detecting the circles, we fill in each circle area to create a binary mask. In this mask, pixel values are set to 1 within the fiducial marker regions and 0 elsewhere, mimicking the format used in our pipeline. This binary mask is then used for further evaluation, enabling a direct comparison of the performance between the 10x pipeline and Vispro.

#### Baseline U-Net

For the baseline U-Net, we follow the design of `pix2pix`^26^, a widely used U-Net variant for image processing tasks. We adopt its U-Net generator as the baseline for marker segmentation and compare its results with the tailored architecture in Vispro. The network generates fiducial marker masks similar to Vispro, enabling a direct comparison of performance.

### Evaluations

#### Fiducial marker identification

To evaluate the accuracy of fiducial marker segmentation, we computed the pixel-level Intersection over Union (IoU) using the manually annotated gold standard as a reference. The IoU is a metric that measures the overlap between the predicted mask and the ground truth mask at the pixel level. It is defined as the ratio of the number of correctly predicted pixels (i.e., the intersection of the predicted and ground truth masks) to the total number of pixels in the union of the predicted and ground truth masks:

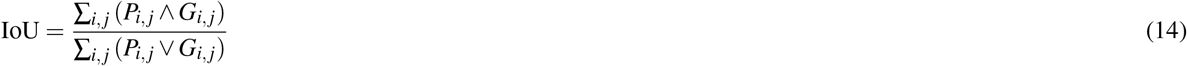

where *P*_*i, j*_ represents the predicted mask at pixel location (*i, j*), *G*_*i, j*_ represents the ground truth mask at pixel location (*i, j*), ∧ denotes the logical AND operation (i.e., the number of pixels where both *P*_*i, j*_ = 1 and *G*_*i, j*_ = 1, representing the intersection), and ∨ denotes the logical OR operation (i.e., the number of pixels where either *P*_*i, j*_ = 1 or *G*_*i, j*_ = 1, representing the union). A higher IoU value indicates a higher segmentation accuracy, as it shows that the predicted mask aligns more closely with the ground truth mask.

All images were divided into 12 groups for 12-fold cross-validation. In each iteration, images from 11 groups were used for training, while images from the remaining group were used for testing.

#### Identifying tissue regions

To assess the accuracy of tissue region segmentation, we manually annotated tissue regions on 30 tissue slices, encompassing a spectrum of difficulty from straightforward to challenging cases. We evaluated two sets of images: the restored images produced by Vispro and the original images from the 10x pipeline. For each image, the tissue probability threshold on the tissue mask 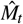 was adjusted to delineate the tissue regions, and the Intersection over Union (IoU) was computed against the corresponding ground truth.

#### Segregating disconnected tissue regions

To evaluate the accuracy of tissue segregation, we manually annotated the disconnected tissue regions on 10 slices, each containing more than one distinct tissue area. We then compared the number of segmented tissue regions produced by Vispro or the 10x pipeline against the gold standard. Additionally, we calculated the IoU for the complete tissue regions compared to the ground truth to account for the impact of tissue segmentation.

#### Cell segmentation

To evaluate cell segmentation performance, we utilized StarDist^34–36^ to detect cells in both the original H&E images and the images processed by Vispro. StarDist is a state-of-the-art segmentation algorithm designed for star-convex object detection in both 2D and 3D images, making it particularly well-suited for accurately segmenting objects with varying shapes and sizes, such as nuclei in histopathological images. We employed the `2D_versatile_he` model, pre-trained for H&E-stained histological images. The main model parameters were configured as follows: `block_size` was set to 4096, `prob_threshold` to 0.01, and `nms_threshold` to 0.001.

For cell segmentation results obtained by each method, we calculated the number of segmented cells falling outside of the manually annotated tissue regions.

#### Image registration

We evaluated the accuracy of image registration in both same-modal and cross-modal settings. For the same-modal setting, we used 20 pairs of Visium images from our collected dataset, where each pair originated from the same study and shared similar textures. For the cross-modal setting, we collected 13 pairs of images from the Visium CytAssist dataset. This dataset involves overlaying images of standard histological glass slides with those processed into Visium slides. While structurally identical, the pairs exhibit slight texture differences, making them ideal for testing cross-modal registration performance.

We employed bUnwarpJ registration^37^, available within the ImageJ/Fiji software^20^, for image registration. bUnwarpJ performs 2D image registration using elastic deformations modeled by B-splines, ensuring invertibility through a consistency constraint. We used the stable version (2.6.13) of the software, configuring the initial deformation to `coarse`, the final deformation to `super fine`, while leaving all other parameters at their default settings.

Two metrics were used for quantitatively evaluations. The first metric is the Structural Similarity Index (SSIM), which measures the similarity between two images by comparing local patterns of pixel intensities, normalized for differences in luminance and contrast. SSIM is particularly sensitive to pixel misalignment, making it a robust metric for evaluating registration performance, especially in the context of same-modal image registration. For the multi-channel data in our case (i.e., RGB with three channels), SSIM is calculated separately for each channel, and the final metric is obtained by averaging the scores across all channels. This approach ensures a comprehensive evaluation of structural similarity for color images. For each channel, the SSIM is mathematically defined as:

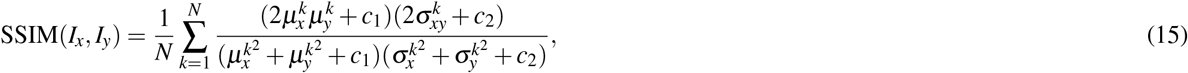

where *I*_*x*_ and *I*_*y*_ are the two input images, *N* is the total number of local image patches. 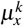 and 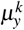 represent the local means, 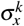 and 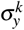 are the local standard deviations, and 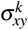 denotes the cross-covariance of the intensity values in the *k*-th patch of *I*_*x*_ and *I*_*y*_. The constants *c*_1_ and *c*_2_ are stability parameters introduced to prevent division by zero.

The second metric is Mutual Information (MI), which quantifies the amount of information shared between two random variables and is widely used in image registration tasks to measure the similarity between images. As MI is calculated from whole-image pixel patterns, it excels at identifying alignment by leveraging structural and contextual consistency rather than relying on simple intensity correspondence, making it particularly effective for estimating registration tasks involving images from different modalities. To handle the continuous intensity values in the image pair *I*_*x*_ and *I*_*y*_, the pixel intensities are first discretized into bins using a 2D histogram. Each bin represents a range of intensity values, and the histogram captures the co-occurrence of intensity values from the two images at corresponding spatial locations. The 2D histogram is then normalized to estimate the joint probability distribution *p*(*x, y*), as well as the marginal probabilities *p*(*x*) and *p*(*y*), where *x* and *y* represent the bins of intensity values from *I*_*x*_ and *I*_*y*_, respectively. These probabilities are then used to compute MI.

Formally, MI is calculated as follows:

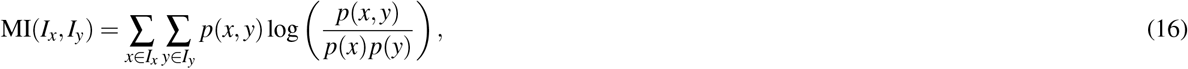

#### Gene imputation

We employed the TESLA software for gene imputation^38^. TESLA leverages the Canny edge detection algorithm from the OpenCV library for contour detection and also offers alternative methods based on the spatial locations of transcripts. We tested all three contour detection methods provided by TESLA and conducted a detailed visual comparison of the imputation outcomes. For consistency, the imputation resolution was set to 50 during implementation.

## Acknowledgments

H.M., A.Z., and Z.J. was supported by the National Institutes of Health (NIH) under award number U54AG075936. Y.Q. and Z.J. was supported by NIH under award number R35GM154865. A.R. was supported by National Science Foundation under award number CAREER-2203741 and NIH under award number R01HL169347.

## Author contributions

H.M., A.Z. and Z.J. conceived the study. H.M. and Y.Q. collected the training data. H.M. conducted the analysis. H.M. and Z.J. wrote the manuscript.

## Code availability

Vispro can be freely accessible at https://github.com/HuifangZJU/Vispro. The Github repository includes detailed instructions for usage and any required dependencies.

