## Supplementary Table S1 and Supplementary Figure S1 for "Vispro improves imaging analysis for Visium spatial transcriptomics"

### Supplementary materials

| Image Resolution | GPU (A4000, 8GB) | CPU |
|---|---|---|
| $\sim$2,000$\times$2,000 | $\sim$5s | $\sim$30s |
| $\sim$15,000$\times$15,000 | $\sim$30s | $\sim$500s |

**Table S1.** Running time of Vispro on GPU and CPU with different image resolutions.

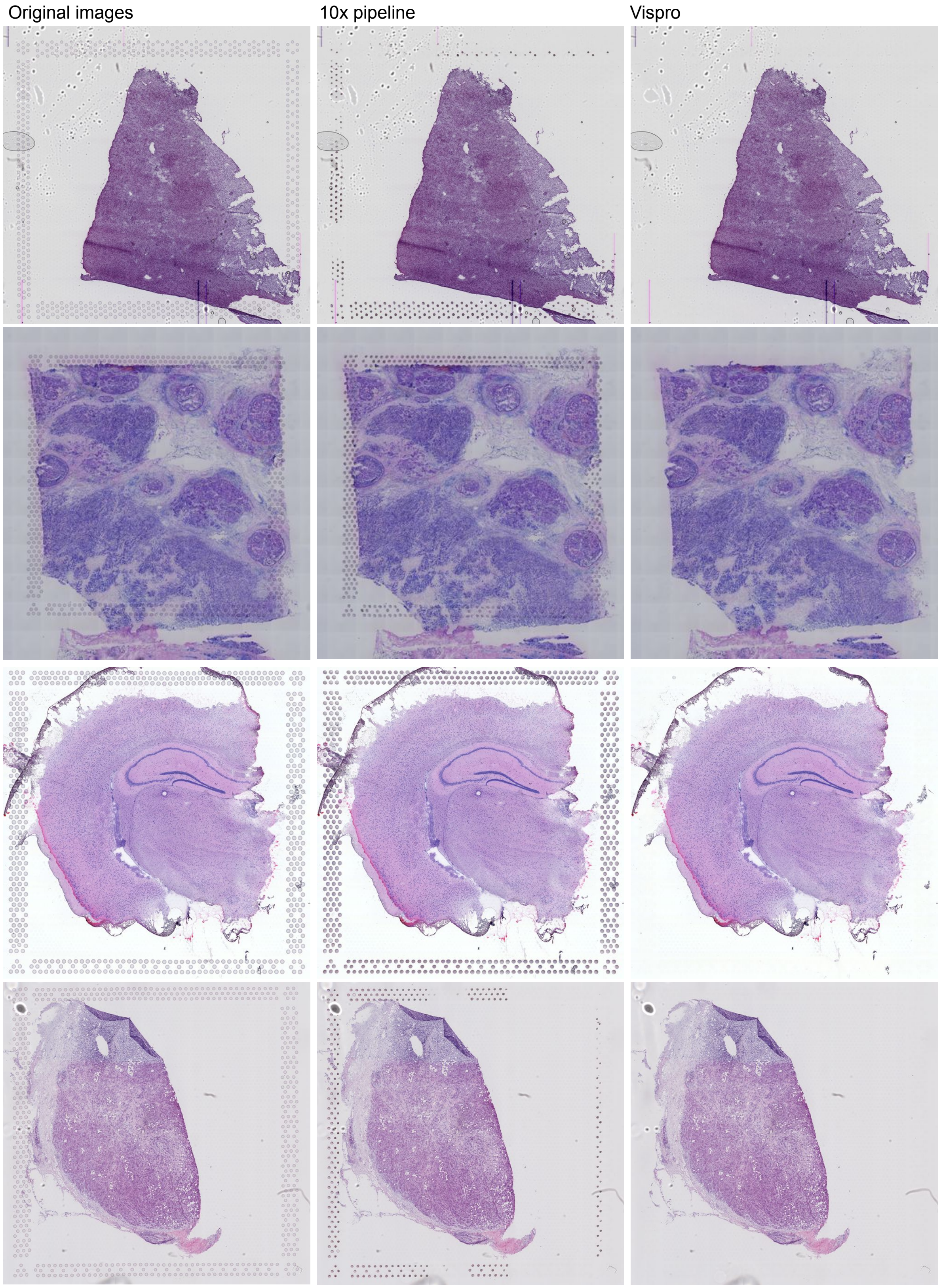


**Figure S1.** Visual comparison of image restoration results. The three columns display, from left to right: the original images, the restored images with fiducial markers identified by the 10x pipeline, and the restored images with fiducial markers identified by Vispro. Each row represents an image from a Visium sample.
